# Experimental evolution reveals a novel avenue to release catabolite repression via mutations in XylR

**DOI:** 10.1101/100602

**Authors:** Christian Sievert, Lizbeth M. Nieves, Larry A. Panyon, Taylor Loeffler, Chandler Morris, Reed A. Cartwright, Xuan Wang

## Abstract

Microbial production of fuels and chemicals from lignocellulosic biomass provides promising bio-renewable alternatives to the conventional petroleum-based products. However, heterogeneous sugar composition of lignocellulosic biomass hinders efficient microbial conversion due to carbon catabolite repression. The most abundant sugar monomers in lignocellulosic biomass materials are glucose and xylose. While industrial *Escherichia coli* strains efficiently utilize glucose, their ability to utilize xylose is often repressed in the presence of glucose. Here we independently evolved three *E. coli* strains from the same ancestor to achieve high efficiency for xylose fermentation. Each evolved strain has a point mutation in a transcriptional activator for xylose catabolic operons, either CRP or XylR, and these mutations are demonstrated to enhance xylose fermentation by allelic replacements. Identified XylR variants (R121C and P363S) have a higher affinity to their DNA binding sites, leading to a xylose catabolic activation independent of catabolite repression control. Upon introducing these amino acid substitutions into the *E. coli* D-lactate producer TG114, 94 % of a glucose-xylose mixture (50 g L^-1^ each) was utilized in mineral salt media that led to a 50 % increase in product titer after 96 h of fermentation. The two amino acid substitutions in XylR enhance xylose utilization and release glucose-induced repression in different *E. coli* hosts, including wild-type, suggesting its potential wide application in industrial *E. coli* biocatalysts.

## Introduction

Microbial biocatalysts such as *Escherichia coli* and *Saccharomyces cerevisiae* have been developed to convert sugars to an array of value-added chemicals, ranging from simple fermentation products to complex terpenoids like artemisinic acid [1]. Lignocellulosic biomass represents a promising renewable feedstock that can support large-scale microbial production processes for fuels and specialty chemicals without interfering with human food supply [2, 3]. Lignocellulose is a complex matrix present in plant cell walls and is mainly composed of polysaccharides and phenolic polymers [2]. D-glucose (the sole monomer of cellulose) and D-xylose (the predominant sugar in hemicellulose) are major sugars found in typical lignocellulosic materials [2, 3]. Although the sugar content in lignocellulosic materials (e.g. agricultural wastes such as corn stover) is higher than 50 % of their dry weight, the heterogeneous nature of lignocellulosic sugars inhibits efficient microbial catabolism and thus decreases production [2, 3]. Industrial microbes such as *S. cerevisiae* and *Zymomonas mobilis* do not natively metabolize xylose and a foreign xylose catabolic pathway must be integrated into these hosts for xylose utilization [4, 5]. Even for bacteria like *E. coli* that natively contain the xylose catabolic pathway, xylose utilization rates and growth rates on xylose are low [6]. More importantly, utilization of xylose is repressed in the presence of glucose due to a global regulatory mechanism called carbon catabolite repression (CCR), a common phenomenon observed in bacteria and fungi, which results in abundant amounts of xylose unused when cells ferment a glucose-xylose mixture [7, 8].

As one of classic global regulatory mechanisms, CCR is well characterized in *E. coli* [7, 8]. The global transcriptional regulator CRP (cAMP receptor protein) plays a central role in modulating transcriptional activation of catabolic operons for secondary sugars such as xylose, arabinose and galactose [7, 8]. The phosphoenolpyruvate:sugar phosphotransferase system (PTS) and the membrane bound enzyme adenylate cyclase (AC; catalyzing the conversion of ATP to cAMP) are also involved in glucose-induced repression of xylose utilization in *E. coli* [7, 8]. The phosphorylation state of EIIA^Glc^, a PTS component encoded by *crr* in *E. coli*, plays a pivotal role in regulating AC activities and cAMP levels according to glucose concentrations [9]. When glucose concentrations are high, the phosphate from EIIA^Glc^ is drained towards the sugars and the dephosphorylated EIIA^Glc^ is not able to activate AC, which results in low levels of cAMP [10]. Without cAMP, CRP cannot activate the transcription of xylose catabolic operons. In contrast, at low glucose concentrations copious amounts of phosphorylated EIIA^Glc^ exist and are able to activate AC and promote cAMP synthesis. If xylose is present, CRP activated by cAMP and the xylose-specific activator XylR (activated when bound by xylose) together co-activate the xylose catabolic operons, *xylAB* and *xylFGH* (Fig. 1*A*) [11]. After xylose is imported by XylFGH, a xylose-specific ATP-binding cassette transporter protein, xylose is converted to xylulose through a reversible one-step reaction catalyzed by xylose isomerase, XylA. Xylulose is then converted to xylulose-5-phosphate by the xylulokinase, XylB, for further degradation via the pentose phosphate pathway and glycolysis (Fig. 1*A*) [12].

**Figure 1:**
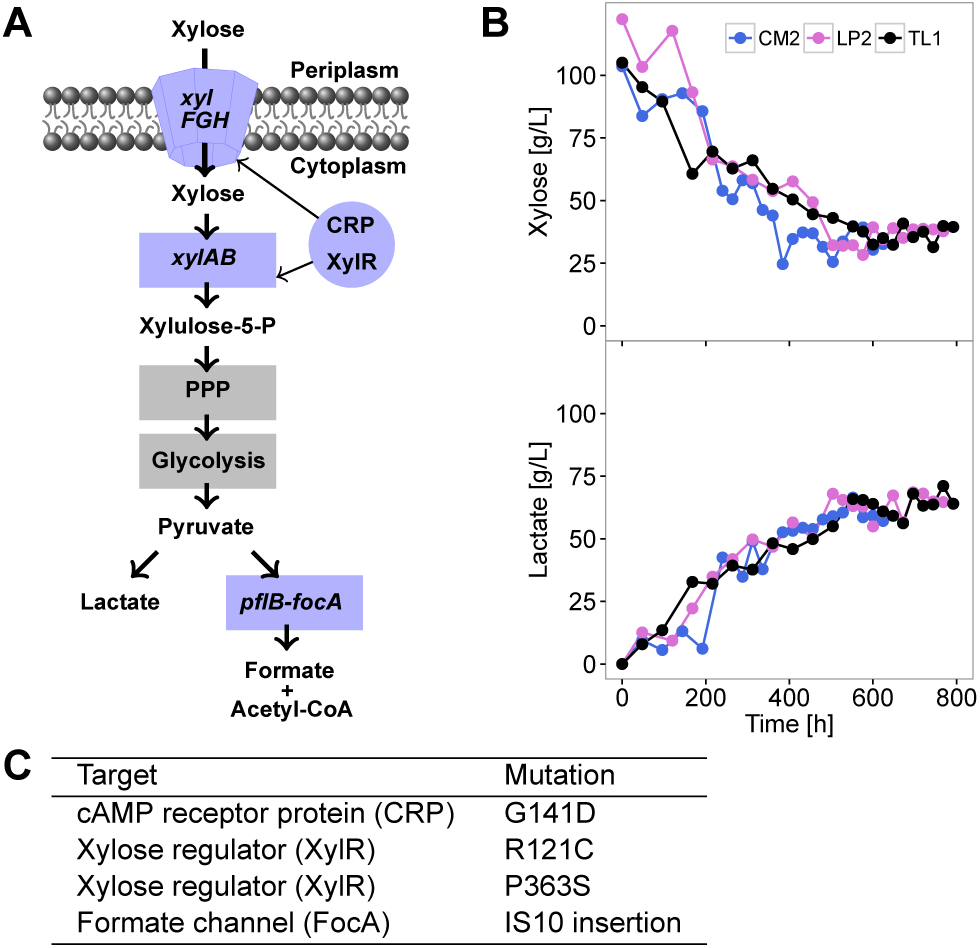
Overview of the adaptive laboratory evolution experiment. (A) Schematic drawing of the xylose to D-lactate pathway. Xylulose-5-P, xylulose-5-phosphate; PPP, pentose phosphate pathway. (B) Serial transfers of *E. coli* strain XW043 were cultured independently three times resulting in three evolved populations, from which CM2, LP2, and TL1 were isolated. Extracellular xylose and D-lactate were determined for each transfer after 48 h of fermentation. (C) Primary convergent mutations were characterized using genome sequencing and allelic replacements.

Releasing CCR by genetic engineering allows microbial biocatalysts to simultaneously utilize glucose and other secondary sugars derived from lignocellulosic biomass and leads to a more efficient fermentation process [7]. Common strategies to engineer sugar co-utilization in *E. coli* include the inactivation of PTS components, such as PtsG and PtsI [13, 14, 15], and mutagenesis of CRP [16, 17]. However, both approaches have caveats and only limited success has been achieved, especially for conditions more relevant to industrial practice such as high sugar concentrations and low-cost media. Inactivation of PTS components also impairs glucose uptake and thus extra efforts to compensate this defectiveness are needed [13, 18]. Theoretically, a cAMP-independent CRP variant should be able to activate the catabolic operons of secondary sugars even in the presence of glucose. However, these cAMP-independent CRP mutants often cannot activate the target operons at the same efficiency as wild-type CRP bound with cAMP [16]. In addition, as an important global regulator, CRP directly regulates the transcriptional expression of more than 400 genes and CRP mutants often have slow growth phenotypes potentially due to unpredictable expressional changes of other important genes [19]. Here, we evolved *E. coli* for enhanced xylose fermentation and identified the convergent genetic basis that increases xylose utilization. By characterizing the transcriptional activation mechanism of xylose catabolic genes, we discovered a simple and effective genetic approach to release CCR in *E. coli*. We identified single nucleotide mutations in *xylR* that increase xylose utilization up to 4-fold in different *E. coli* strains when fermented in a glucose-xylose mixture (50 g L^−1^ of each sugar). This discovery has the potential to enable different *E. coli* biocatalysts to simultaneously convert major sugars from lignocellulosic biomass into value-added chemicals.

## Results

### Identification of primary genetic changes of *E. coli* adaptation for improved xylose fermentation to D-lactate

We hypothesized that characterization of repeated evolutionary trajectories would reveal the convergent causative mutations that improve xylose fermentation. A previously engineered D-lactate producing strain XW043 [20] was independently evolved three times in mineral salt media containing 100 g L^−1^ xylose (Fig. 1*B*). The bacterial population was maintained at the exponential or early stationary phase in a fermentation vessel by serially transferring cultures into new media as previously described [20, 21]. In all three evolutionary trajectories, a rapid adaptation occurred that simultaneously increased xylose catabolism, lactate titer and cell growth (Fig. 1*B*, S1). Since lactate production is the only fermentation pathway supporting cell growth in XW043 under oxygen-limiting experimental conditions [20], increased xylose catabolism would lead to higher cell growth. At the end of three evolution experiments, there was an approximately 5-fold increase of lactate titers at 48 h compared to the ancestor strain XW043, accompanied with an increase in yield from 0.6 to 0.8–0.9 g lactate per gram xylose (Fig. 1*B*). To understand the genetic changes responsible for the increased xylose catabolism, we sequenced the genomes of the ancestor XW043 and three representative evolved clones (one from each evolved population), designated as strains TL1, CM2, and LP2 using Illumina paired-end sequencing. Each clone was sequenced twice with an average 24-fold coverage per library. By applying a comprehensive analysis pipeline (Details in SI Methods), overall 5 point mutations, a 1-bp deletion, 15 duplications and 7 deletions in mostly uncharacterized proteins were found in three evolved isolates compared to XW043 (Table S2). Additionally, we detected 60 breakpoints as an indicator for structural rearrangements occurring in all three evolved clones (Table S2). Of all the mutations detected, three point mutations, one per clone, occurred at the transcriptional co-activators CRP and XylR that are critical for xylose catabolism (Fig. 1*A*), suggesting a potential result of convergent evolution to relieve the predominant metabolic constraints in the ancestor strain. The three independently isolated point mutations result in amino acid changes in CRP (G141D in LP2) and XylR (R121C in CM2 and P363S in TL1). In addition, an IS10 insertion was identified in *focA* reading frame for all three evolved clones at different positions (Fig. 1*C*, S4*C*), thereby suggesting an independent origin for each insertion and serving as a strong indicator of the potential benefit of *focA* inactivation for xylose to lactate fermentative production.

### Characterization of physiological effects of convergent mutations

We hypothesized that the identified convergent mutations (*focA* inactivation and missense mutations in *crp* and *xylR*) are primary genetic changes that improved xylose catabolism and lactate fermentation. Genetic mechanisms of evolved phenotypes can be potentially explained by allelic replacement, in which the wild-type copy of the ancestor is replaced by a mutant copy in the descendant, or vice versa [22, 23]. Introduction of a single point mutation *crp* (G141D) in XW043 background doubled both xylose utilization and D-lactate titer after 96 h fermentation, and growth rate and final biomass were also increased (Fig. 2*A*, S2). A 32 kb chromosomal region containing *xylR* and other xylose catabolic genes is duplicated in the ancestor XW043 and all evolved strains as indicated by genome sequencing (Fig. S3) probably because the precursor strain of XW043 was adapted for using hemicellulose hydrolysates as the carbon source [20]. Only one of the two *xylR* copies was mutated (R121C in CM2 and P363S in TL1), thereby conferring a quasi-heterozygous genotype (Fig. S3). We were not able to replace *xylR* with its mutant alleles in XW043 using λ-red recombinase-mediated homologous recombination presumably because the second copy of wild-type *xylR* tended to replace the selection marker due to a favored intrachromosomal recombination. Instead, using λ-red recombinase-mediated intrachromosomal recombination approach (Details in SI Methods), we replaced the *xylR* mutations with a wild-type copy in both CM2 and TL1 strains. This chromosomal modification to restore the wild-type allele decreased xylose utilization rates of the initial 24 h by 40 % and 65 % compared to CM2 and TL1, respectively (Fig. 2*B*, *C*). Accordingly, the modified strains had decreased lactate production and cell growth in both CM2 and TL1 backgrounds (Fig. 2*B*, *C*, S2). However, restoring the *xylR* mutations in CM2 or TL1 back to wild-type did not produce a complete ancestral phenotype in terms of fermentative growth and xylose utilization (Fig. 2*B*, *C*), suggesting the presence of other beneficial mutations important for enhanced xylose utilization.

**Figure 2:**
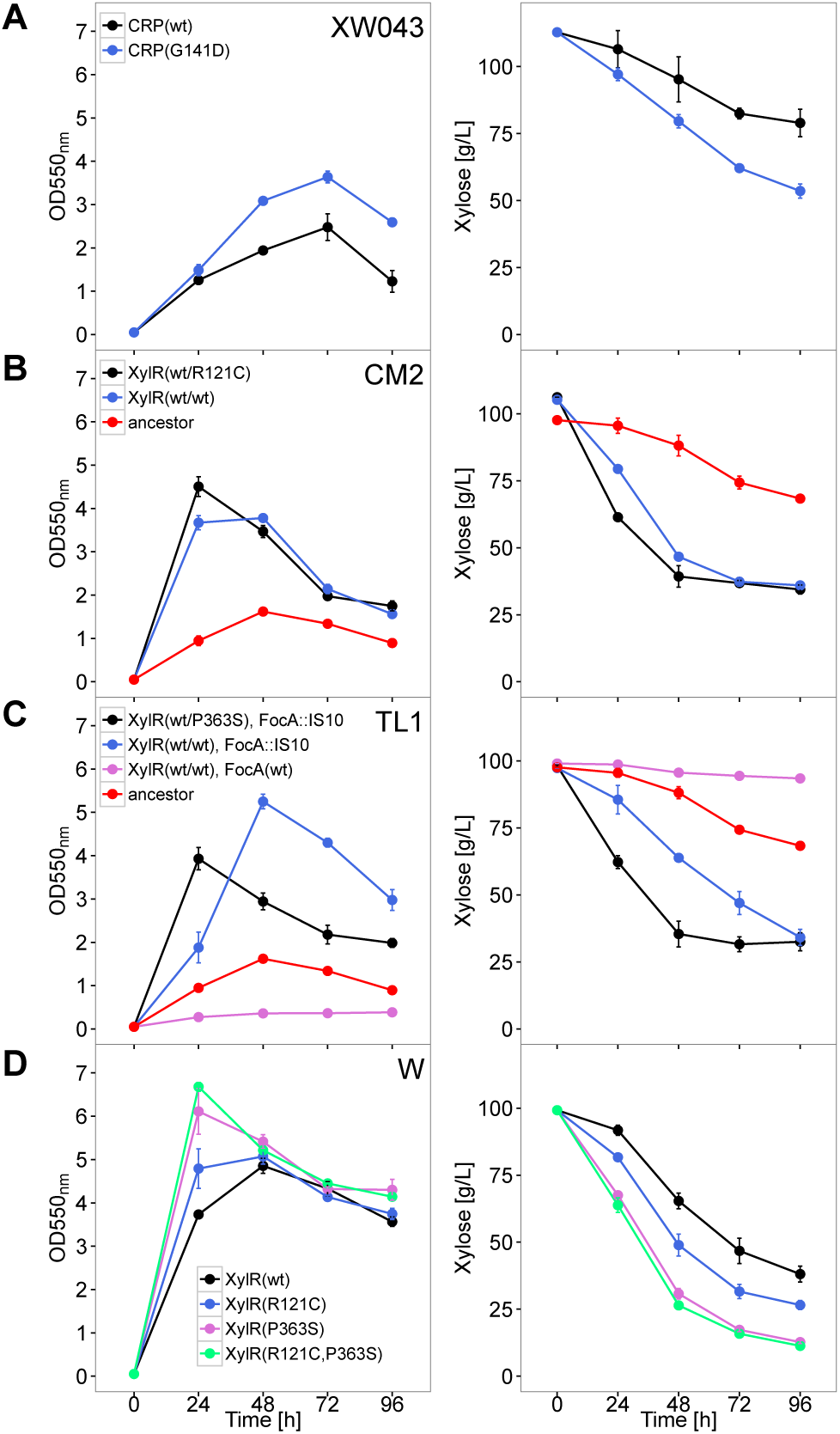
Evaluation of *crp, focA*, and *xylR* mutations. Through allelic replacements, the genotypes of appropriate genes were modified in different genetic backgrounds: the ancestral XW043 (A), the evolved CM2 (B) and TL1 (C), and the wild-type strain W (D). Fermentation performance was determined by measuring the cell optical density (left) and extracellular xylose concentration (right) over 96 h. Point mutations are indicated by the relevant amino-acid substitution. Error bars represent SEM with *n* = 3. wt, wild-type.

Insertion of *focA* by IS10 is another convergent event for all three evolved isolates, suggesting a potential benefit for xylose to lactate conversion (Fig. 1). FocA is a bidirectional formate transporter that regulates intracellular formate levels [24]. Since IS10 elements are repetitive in the genome, we were not able to reconstruct the identical IS10 insertion in the ancestor background. However, an in-frame deletion of *focA* in the ancestor only showed a marginal positive effect on xylose fermentation (Fig. S4*A*), thereby indicating the presence of a more complex mechanism besides *focA* inactivation alone to cause enhanced xylose utilization. The IS10 insertions in *focA* had negative polar effects on transcription of the downstream gene *pflB* as shown by quantitative reverse-transcription PCR (qRT-PCR) in all evolved clones (Fig. S4*B*). The *pflB* gene encodes pyruvate formate-lyase which competes against the lactate production pathway for the common substrate pyruvate with a lower *K_m_* value than lactate dehydrogenase (Fig. 1*A*) [24, 25]. Decreased expression of *pflB* may help redirect carbon flow to lactate dehydrogenase. To test the epistatic interaction between the identified convergent mutations, we restored wild-type *focA* and *xylR* in evolved background TL1 to reconstruct an ancestor phenotype (Fig. 2*C*). Interestingly, this strain (CS4: TL1 *xylR* wt/wt, *focA* wt) performed even worse than XW043, indicating that both the *xylR* mutation and *focA* IS10 insertion are primary causative mutations with a potential synergy effect and that there is possibly one or further mutations which epistatically interact with the described mutations.

To test if the identified *xylR* mutations have a universal effect and if the heterozygous *xylR* copies are required to improve xylose utilization, we substituted wild-type *xylR* with its mutant version (R121C, P363S, or both) in wild-type *E. coli* W (ATCC9637) (Fig. 2*D*). All mutants increased xylose utilization and cellular fermentative growth to different degrees (Fig. 2*D*). Xylose utilization rates were increased 2.4 and 4.3-fold compared to wild-type strain within the initial 24 h for *xylR* R121C and P363S substitutions, respectively (Fig. 2*D*). We next combined both *xylR* mutations in a wild-type strain and observed a slightly additive effect (Fig. 2*D*). At the end of 96 h fermentation, 40 g L^−1^ xylose remained unused in the broth for wild-type strain while only 10 g L^−1^ for *xylR* P363S substitution or *xylR* R121C and P363S substitution (Fig. 2*D*).

### XylR variants show enhanced activity in vitro

CRP G141D was previously identified as a mutant with an altered allosteric mechanism and position 141 is involved in the interfacial interactions between subunits and domains [16, 26]. Potentially due to the CRP G141D mutation in the evolved strain LP2, the transcription of xylose catabolic genes was upregulated by more than 10-fold as measured by qRT-PCR (Fig. S5). Unlike CRP and its mutant variants, which are well characterized, much less is known about XylR. We examined the effects that *xylR* mutations potentially had on the transcriptional levels for its responsive genes which are organized as two operons *xylAB* and *xylFGHR* (Fig. 3*A*). The *xylAB* and *xylFGH* transcripts were increased at least 20-fold in CM2 with *xylR* (wt/R121C) and 10-fold in TL1 with *xylR* (wt/P363S) compared to the ancestor strain XW043 (Fig. 3*B*). The *xylR* transcript alone shows only between 3.1- and 3.6-fold upregulation suggesting that the *xylFGHR* transcript is partially degraded at the 3’ end or read-through of the RNA polymerase is prevented by clashing with proteins bound at the *xylR* specific promoter. Since the *crp* transcript level is not significantly changed (Fig. 3*B*), we concluded that the XylR mutations are the primary reason for upregulation of the *xyl* operons.

**Figure 3:**
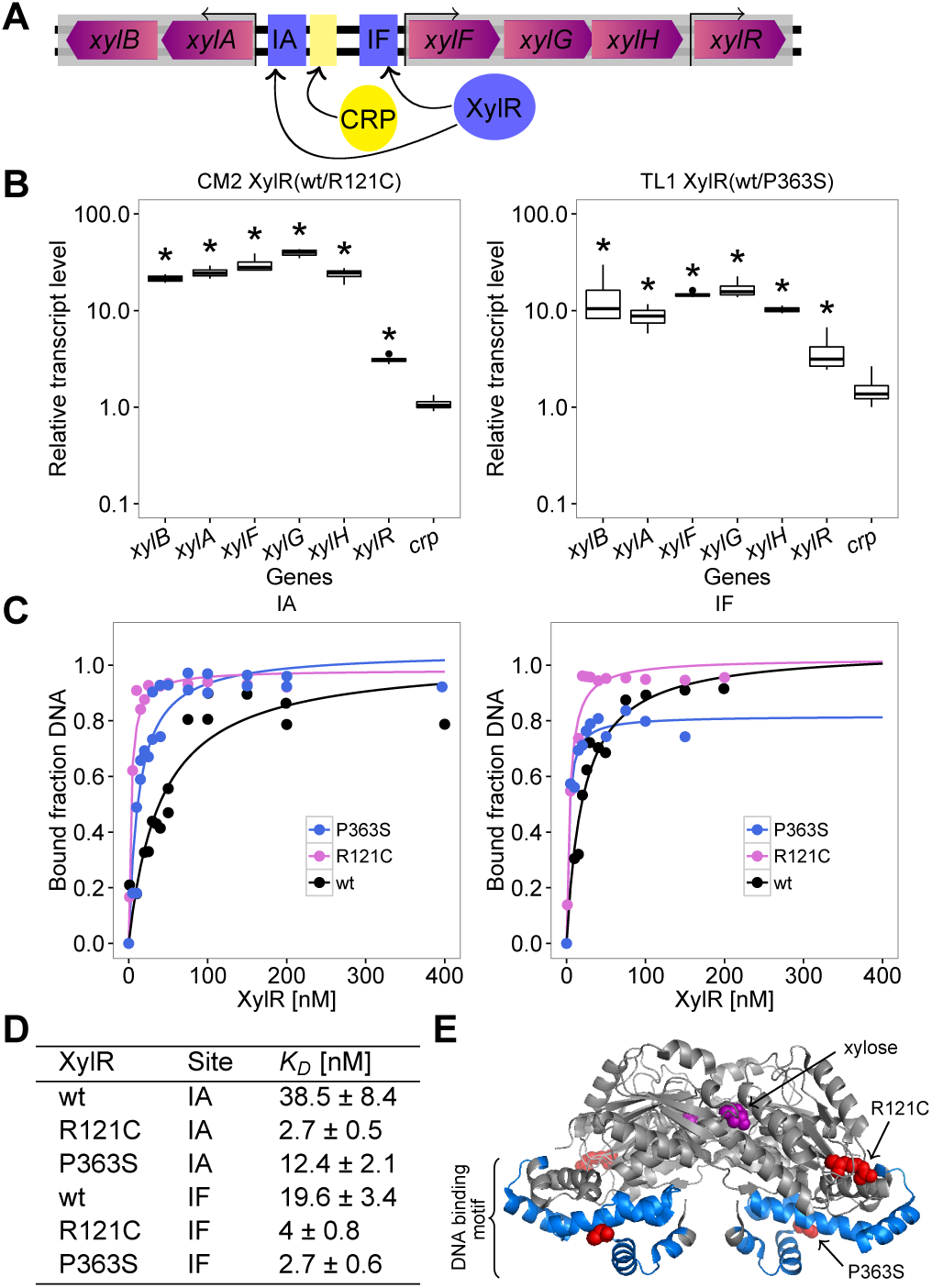
Molecular mechanism of enhanced xylose utilization by XylR mutations. (A) Schematic drawing of the *xyl* operons (gray box) including their transcription start point (arrow). Binding sites of CRP and XylR are indicated as yellow and blue boxes. XylR has two known binding sites IA and IF. The given size does not reflect real proportions. (B) Relative transcript quantification of selected genes of strains CM2 and TL1 with XylR variants R121C or P363S, respectively. The data indicate the fold change of transcript level in relation to the ancestor XW043 background and is normalized by the transcript level of 16S ribosomal RNA (*rrsA*), *n* = 4, unpaired Student’s t-test indicates significance at *p* < 0.05 (*). (C) Fitted data and (D) the determined dissociation constant (*K_D_*) from the electrophoretic mobility shift assays to determine the binding affinity of different XylR variants with their known binding sites IA and IF, respectively. (E) Modeled structure of the dimeric wild-type XylR [27]. Following features are highlighted: xylose (purple), mutated residues (red), and DNA binding domain (blue).

To further study the molecular mechanism causing enhanced transcriptional activation by XylR variants, we conducted electrophoretic mobility shift assays using three purified N-terminal His-tagged XylR variants (wild-type, R121C and P363S) and DNA fragments containing the binding sites IA and IF located in the intergenic region between *xylAB* and *xylFGH* (Fig. 3*A*) [27]. The dissociation constant *K_D_* for each site was determined to measure their binding affinity (Fig. 3*C*, *D*, S6*A*, *B*). Although the gel based assay allows for only a rough approximation, our data for the wild-type XylR is consistent with a previous report showing a *K_D_* of 33±0.8 nM for IA and 25±0.6 nM for IF, respectively (Fig. 3*D*) [27]. The evolved XylRs have a significantly lower *K_D_* ranging from 3- to 14-fold depending on which binding site and variant (Fig. 3*D*, *E*), suggesting that a higher binding affinity leads to a more stable transcription initiation complex and a subsequently higher transcriptional rate. Moreover, wild-type XylR could not bind operator sequences in the absence of xylose since the majority of DNA fragments remained unbound in the gel (Fig. S6*C*). In contrast, both XylR variants bound DNA even without xylose (Fig. S6*C*), suggesting that these XylR variants have xylose-independent activities. A two-fold increase in binding of wild-type XylR to its responsive site was observed when 100 μM xylose was added, but similar enhancement was much less significant for both XylR variants (Fig. S6*C*).

### Amino acid substitutions in XylR release CCR

The higher binding affinity of the evolved XylR and the concomitant xylose independency led to the hypothesis that these amino acid substitutions in XylR are able to release CCR. Wild-type *E. coli* W was used as the test strain for different genetic modifications to release CCR. Batch fermentations were conducted using a mineral salt medium AM1 containing 100 g L^-1^ a glucose-xylose mixture (50 g L^−1^ each). The wild-type strain consumed all glucose but only 16 % of the xylose after 96 h of fermentation (Fig. 4*A*, S7). In comparison, the XylR variants R121C or P363S released CCR, and glucose and xylose were consumed simultaneously, eventually leading to 61-69 % of the xylose utilized (Fig. 4*A*, S7). LN6, the strain with both R121C and P363S substitutions, utilized 87 g L^−1^ of the total sugars compared to 57 g L^−1^ for wild-type (Fig. 4*A*). The xylose utilization rate was increased 4-fold compared to wild-type (81 % of the xylose used after 96 h fermentation) and only 3 g L^−1^ glucose remained unused in the fermentation broth (Fig. 4*A*, S7). CRP G141D alone or together with *xylR* mutations did not enhance sugar co-utilization (Fig. 4*A*). Investigation of known mutations that relieve CCR such as Δ*mgsA* [28] and *crp*\* (I112L, T127I and A144T) [29, 30] was also conducted in the same wild-type background. CRP* only increased xylose utilization to 37 % with impaired cell growth while the *mgsA* deletion had no benefit in wild-type background (Fig. 4*A*, S7).

**Figure 4:**
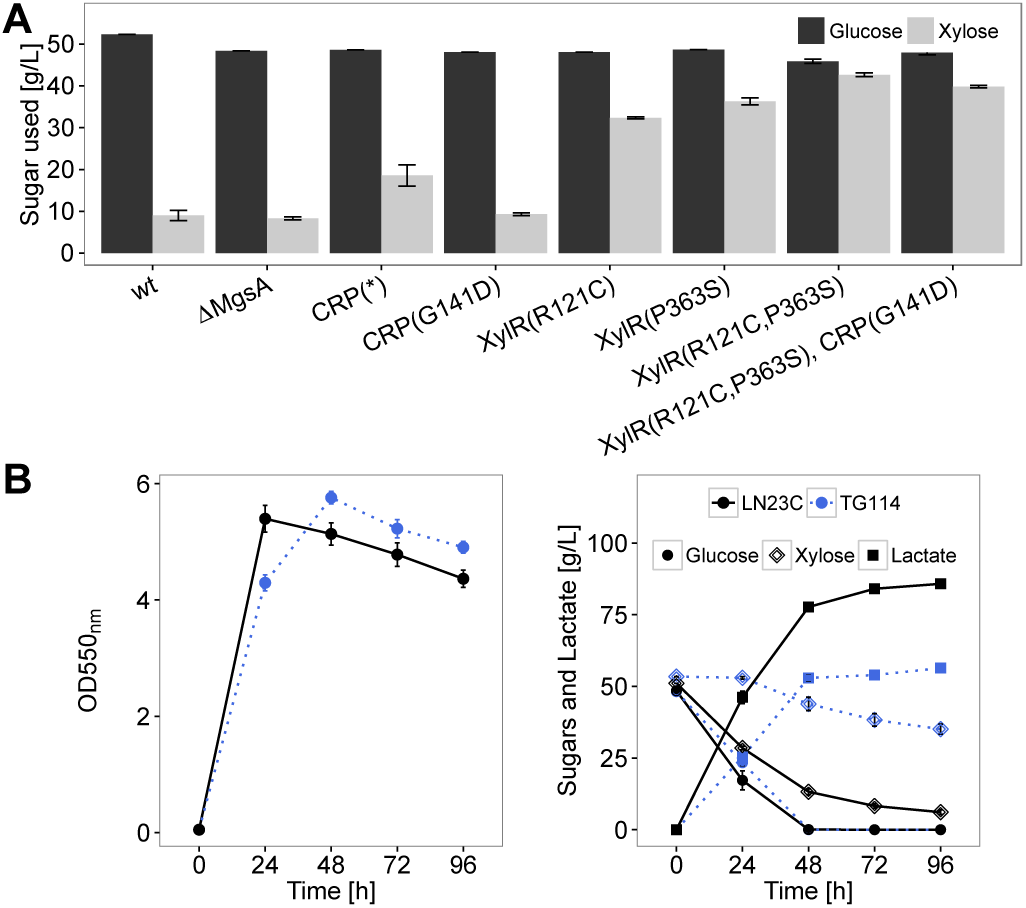
Co-utilization of glucose-xylose mixtures by batch fermentations. (A) *E. coli* W was engineered with different genotypes including previously reported genetic traits such as *mgsA* deletion (ΔMgsA) and CRP*, and identified CRP and XylR variants to compare their effect of releasing CCR. Batch Fermentations were performed using glucose-xylose mixtures (50 g L^−1^ of each sugar) for 96 h. wt, wild-type. (B) The fermentation of TG114 and its modified strain LN23C with the R121C/P363S substitution in XylR using sugar mixtures. Cell optical density (OD550_nm_), xylose, and D-lactate concentrations were determined. Error bars represent SEM, *n* = 3.

To further demonstrate the application of our discovery in converting sugar mixtures into renewable chemicals, we substituted wild-type *xylR* with *xylR* (R121C and P363S) of a previously engineered D-lactate *E. coli* producer TG114, which efficiently converts glucose into D-lactate [21]. Fermenting TG114 in a glucose-xylose mixture, 50 g L^-1^ of each sugar, showed that only 34 % of the xylose was used due to CCR (Fig. 4*B*). Introducing R121C and P363S substitutions in *xylR* enabled the modified strain to use 88 % of the xylose while all glucose was consumed within 96 h. Consequently, D-lactate production increased to 1.5-fold compared to TG114 with a final titer at 86 g L^−1^ and a yield of 0.91 g g^−1^ sugars.

## Discussion

We characterized the primary genetic changes of adaptation for xylose utilization in an *E. coli* D-lactate producer and found a novel way to effectively release CCR in *E. coli*. By deleting competing fermentation pathways in the ancestor strain, the possible evolutionary trajectories were strictly constrained to xylose-lactate conversion and convergent mutation patterns were observed in three parallel evolved populations. In contrast to the convergent mutations in *crp* and *xylR* identified in this work, previous research using similar experimental evolutionary approaches showed that beneficial mutations for enhanced xylose fermentations in *E. coli* biocatalysts occurred at sugar transporter genes, such as *galP* and *gatC* [31, 32]. This suggests that there are potentially multiple evolutionary solutions to the same problem and bacterial genotypic backgrounds may predetermine evolutionary trajectories. Current high-throughput next-generation sequencing capacity has significantly outpaced reverse engineering processes to characterize the causative mutations. One time-consuming bottleneck is to distinguish the causative mutations from mutational noise and sequencing errors [22]. Here, two strategies were employed to effectively identify most critical mutations for the adapted phenotype. First, characterization of multiple independent evolutionary trajectories from the same ancestor revealed the convergent mutations important for adaptation to experimental conditions as demonstrated in this work and other laboratory evolution research [33]. Second, as sequencing technical errors were excluded by sequencing each genome twice with high coverage, linkage disequilibrium is probably the only source of noise which we further reduced by focusing on only an individual evolved clone instead of the whole bacterial population.

We identified a G141D substitution in CRP enhancing transcription of *xyl* operons in *E. coli*. This mutation was previously reported to alter allosteric regulation [16]. Position 141 is part of a hinge important for the intramolecular transduction of the activation signal [34]. CRP G141D did not release CCR (Fig. 4*A*), suggesting that this variant remains to be responsive to cAMP, which is consistent with other previously reported mutations at this position [26]. We hypothesize that a polar residue Asp at the position 141 orients the DNA binding domain for a better interaction with operator sequences resulting in an increased expression compared to wild-type. Interestingly, an intrinsically active CRP homolog of the plant pathogen *Xanthomonas campestris* has an Asp residue at the corresponding position to G141 of *E. coli* CRP. A substitution of this Asp residue with the nonpolar residue Ala reduces the binding affinity to its promoter by approximately 12-fold [35]. In contrast to CRP G141D, the identified XylR variants not only enhance xylose utilization but also release CCR leading to an efficient co-utilization of glucose-xylose mixtures (Fig. 4). The AraC-type transcription factor XylR has two helix-turn-helix (HTH) motifs and its N-terminal ligand binding domain contains a unique periplasmic-binding protein fold that is structurally related to LacI/GalR transcription factors [27]. Upon xylose binding, XylR undergoes a conformational change orienting the two HTH motifs of the DNA binding domain in an active conformation [27]. In one instance, a Pro residue of XylR was substituted with Ser (P363S) at the site that connects the two HTH motifs likely causing a significant reorientation and tighter binding to the DNA (Fig. 3*E*). The second variant is a R121C substitution which is neither in close proximity to the DNA nor to the ligand binding domain (Fig. 3*E*). The region is also not known to participate in signal communication from the N-terminal ligand binding to the C-terminal DNA binding domain [27]. However, the effect on the binding affinity is similarly increased in both variants. A possible mode of action might be an interaction of residue 121 with Thr185 which is directly connected to a helix motif that promotes dimerization of XylR. Cys is less bulky than Arg which might stabilize dimerization due to a reduced steric hindrance.

CCR in *E. coli* can be dramatically released simply by amino acid substitution in XylR as demonstrated here (Fig. 4). The likely mechanism is that the enhanced binding of XylR variants to the operator sequences between *xylAB* and *xylFGH* operons enables independency from both CRP and xylose, thereby resulting in the observed sugar co-utilization. Multiple genetic engineering approaches have previously been developed to release CCR in *E. coli*. Inactivation of *mgsA* significantly enhanced sugar co-utilization in an ethanologenic *E. coli* [28]. However, glucose-induced repression is still severe in two tested strains with inactivated *mgsA*, LN24B (*E. coli* W Δ *AmgsA*) and TG114 (*mgsA* is deleted to prevent product impurity) (Fig. 4), suggesting its limited application in a broad range of *E. coli* catalysts. Several cAMP-independent CRP mutants were discovered to release CCR to some extent [16, 36]. But cell growth is often stunted by *crp* mutations and sugar coutilization is not efficient (Fig. 4*A*, S7). Disruption of the PTS system (deletion of *ptsG* or *ptsl*) is an effective approach to release CCR, but glucose uptake is impaired and need to be re-engineered for glucose utilization, which involves extra genetic modifications or adaptation [7, 13, 18]. In this study, by simply substituting two residues in XylR of an established D-lactate producer, this modified strain is able to simultaneously ferment 50 g L^−1^ glucose and 43 g L^−1^ xylose and produce 86 g L^−1^ lactate within 96 h in a mineral salt medium (Fig. 4*B*), which shows advantages compared to previously engineered strains for lactate fermentative production using sugar mixtures in terms of the xylose utilization rate, lactate yield and titer under a similar batch fermentation condition [14, 37]. This genetic approach may be effective for other *E. coli* biocatalysts since these XylR variants have a similar positive effect in a wild-type strain (Fig. 4*A*).

The ratio between glucose and xylose in many lignocellulosic hydrolysates is usually higher than that was used in this study (1:1 by weight) [2] so that the positive effect of XylR substitution on co-utilization of sugars derived from lignocellulosic hydrolysates is expected to be even greater due to the relative lower ratio of xylose. Moreover, these XylR variants may also reduce arabinose-induced repression which is caused by a competitive binding of AraC (activated by L-arabinose) to the regulatory regions of *xyl* operons [38]. Enhanced binding of these XylR variants to the operator sequences will potentially compete against AraC and achieve co-utilization of arabinose and xylose.

## Materials and Methods

Detailed description of the materials and methods can be found in SI Methods. The used strains, plasmids, and primers are summarized in Table S1. Chromosomal modification was conducted using two-step λ-red recombinase-mediated homologous recombination as previously described [39]. For batch fermentations and adaptive evolution experiments, *E. coli* was grown at 37 °C in a pH-controlled fermentation vessel using AM1 mineral salt media containing a defined carbon source [20, 21]. Genomes were sequenced using a MiSeq Illumina sequencer generating 600 nt paired-end reads and all sequencing reads were deposited in NCBI (SRA accession: SRP083931).

## Acknowledgments

We thank Scott Bingham, David Winter, and Kael Dai for technical assistance. This work was supported by Arizona State University and NIH grant R01-HG007178.

## SI Methods

### Strains, Plasmids and Growth Conditions

Strains used in this study are listed in Table S1. The D-lactate producing *E. coli* XW043 and TG114 (a derivative of *E. coli* W, ATCC9637) were kindly provided by Dr. Lonnie Ingram’s lab at the University of Florida [20, 21]. Strains CS1, CS2, CS3, CS4, CS5, LN4, LN5, LN6, LN8, LN24B, LN21, LN8C, and LN23C were genetically engineered in this study. PC05 containing CRP* was previously constructed [30] and generously provided by Dr. Patrick Cirino at University of Houston.

Plasmid pXW1 was constructed by inserting a native sequence (104 nucleotides after *sacB* ORF) containing a complete terminator sequence into a previously made plasmid pLOI4162 [39]. The *sacB* with its complete terminator serves as an effective counter-selection marker for chromosomal engineering. The original plasmid pLOI4162 containing the *cat-sacB* cassette was kindly provided by Dr. Lonnie Ingram at the University of Florida. The *xylR* ORF and its variant genes were cloned between *Nde*I and BamHI sites in the pET-15b for expression and purification. To create pXylRwt (wild-type *xylR* cloned into pET-15b) the wild-type *xylR* was amplified from genomic DNA in the ancestral strain XW043 by PCR using the Phusion High-Fidelity DNA polymerase (New England Biolabs) with the *Bam*HI site at the end of *xylR* ORF. The pET-15b was digested with the *Nde*I and *Bam*HI restriction enzymes and purified before ligation. The purified wild-type *xylR* PCR product was ligated into the *Bam*HI site of pET-15b at 16 °C for 1 h using the Quick T4 DNA ligase from New England Biolabs. DNA ends were filled by the DNA Klenow fragment from Thermo Scientific by incubating for 15 min at room temperature then at 16 for 3 h to self-ligate and create a complete plasmid. This plasmid was then transformed into TOP10 competent *E. coli* cells. Sequencing results showed that 9 additional nucleotides accidentally were created at the cloning site downstream of the *xylR* start codon in the selected positive transformant. Quick-change PCR was used to remove these additional nucleotides and sequence of *xylR* ORF was confirmed correctly by Sanger sequencing. Similarly, Quick-change PCR was used to modify pXylRwt to create plasmids containing *xylR* variants, pXylR R121C and pXylR P363S. Primers containing the desired mutation were used to amplify the plasmid DNA using the Phusion High-Fidelity DNA polymerase in 12 PCR cycles. After removing the template plasmid DNA by *DpnI* (New England Biolabs) digestion the newly synthesized DNA was transformed into TOP10 cells and positive constructs were sequenced to confirm the desired mutation. The cloning strain TOP10 and pET-15b were kindly provided by Dr. David Nielsen at Arizona State University.

During strain construction, cultures were grown aerobically in shake flasks containing Luria broth. Ampicillin (50 mg L^−1^), kanamycin (50 mg L^−1^), or chloramphenicol (40 mg L^−1^) was added as appropriate. All fermentations were performed using AM1 mineral salt medium supplemented with different carbon sources [20]. Prior to fermentations, strains were streaked on AM1 agar plates with 2 % xylose (w/v) from −80 °C glycerol stocks and incubated at 37 °C for 20 h in a container filled with argon gas. Cells from plates were then grown in AM1 medium containing 2 % xylose (w/v) buffered with 100 mM MOPS in Erlenmeyer flasks as seed cultures. Seed cultures were incubated at 37 °C for 16 h before being harvested by centrifugation (5 min, 6750 g, 4) and used as a starting inoculum for batch fermentations. Batch fermentations were carried out in 500-mL fermentation vessels containing 300 mL AM1 media (100 g L^−1^ xylose or varied sugar mixtures). The initial cell density was 0.022 mg dry cell weight (dcw) mL^−1^, with an optical density (OD) value of 0.05 as measured by a Beckman DU®730 spectrophotometer at 550 nm. Fermentations were maintained at pH 7.0 by the automatic addition of 6 M KOH as previously described [21]. Glucose, xylose, and other organic acids in fermentation broth were measured by high-performance liquid chromatography (HPLC) using an Aminex^®^ HPX-87H column (Bio-Rad) and 4 mM sulfuric acid as the mobile phase [21].

### Laboratory Evolution Experiments

Strain XW043 was consecutively transferred into fresh media in the above mentioned fermentation vessels at approximately 24–48 h time periods for a total of 20 transfers (~150 generations). 0.022 mg dcw mL^−1^ (0.05 as OD550) and 0.0044 mg dcw mL^−1^ (0.01 as OD550) were used as the starting cell densities for early transfers and late transfers (after the 5th transfer), respectively. Approximately every two transfers, a sample of the bacterial population was stored at −80 °C. XW043 was evolved independently three times to improve its xylose utilization ability. Three or more independent clones were selected from each evolved population (the 20th transfer), and their fermentation performance was universally similar to that of the evolved populations. One representative clone for each evolved population was selected for further genome sequencing. These three clones are designated as TL1, CM2, and LP2 strains.

### Genetic Methods

Red recombinase technology was employed to perform seamless chromosomal deletion, gene replacement, and intra-chromosomal recombination as previously described [39, 40]. Plasmids and primers used in this study are listed in Table S1. During strain constructions, 5 % (w/v) arabinose and 10 % (w/v) sucrose were used for λ Red recombinase induction and counter-selection of *sacB* negative strains, respectively. The positive clones from genetic manipulation were verified by colony PCR and Sanger sequencing.

### Genome Sequencing, Assembly and Identification of de novo Mutations

Genomic DNA from strains XW043, TL1, LP2, and CM2 was extracted and purified using the Promega Wizard genomic DNA purification kit. Extracted DNA was sequenced using the MiSeq Illumina sequencer by the DNASU Next Generation Sequencing Core at Arizona State University. DNA was fragmented to an average size of 500 bp and adapter-ligated fragments were sequenced in both directions with coverage of at least 20×. Each strain was sequenced twice to minimize technical errors, and all sequencing reads were deposited in NCBI (SRA accession: SRP083931). For SNP identification, the best practices work flow for variant detection developed by the Broad Institute was used. Briefly, the reads were mapped with Bowtie2 against the closest relative with a sequenced genome, LY180 (accession number CP006584). Duplicate reads were marked using Picard Tools. The alignment was adjusted using GATK’s function for indel realignment and base recalibration. For the latter, the realignment was also analyzed with the mpileup function of SAMtools to call a set of putative variants which was then used to recalibrate. From this final alignment, the raw variants were called using the UnifiedGenotyper from GATK and annotated using SnpEff. Larger duplications or deletions were identified by analyzing the read coverage with a sliding window approach using CNVnator and a bin size of 50 bp. Other types of structural rearrangement were identified using a de novo assembly approach. The assembled contigs by Velvet were combined using MAUVE and compared against the ancestral data using MUMmer. All variants were filtered using SnpSift for occurrence in both sequencing duplicates and absence in the ancestral strain. Structural variants from de novo assembly were only called if present in all three parallel evolved populations in order to reduce the false-positive rate and to identify only most significant changes.

### Quantitative Real Time PCR

Total RNAs from 0.22 mg dcw (~ 5 × 10^8^ cells) were extracted using the RNeasy mini kit from Qiagen. Cells were broken in the lysis buffer from the extraction kit using a FastPrep^®^-24 disrupter (MP biomedicals, Solon, OH) with maximum speed twice (1 min for each time). An optional on-column DNase treatment was performed. Isolated RNA was evaluated by agarose gel electrophoresis. For reverse transcription, the Maxima Reverse Transcriptase from Thermo Scientific was used and 500 ng total RNA was primed with random hexamer oligonucleotides. 2 μL of a 1:20 cDNA dilution was used in subsequent Real Time PCRs conducted with the Maxima SYBR Green/ROX qPCR Master Mix from Thermo Scientific. Relative transcript levels were determined using the comparative C_T_ method [41]. The data was normalized with the level of 16S ribosomal RNA (*rrsA*). Unpaired Student’s t-test indicates significance at *p* < 0.05 (*) with n = 4. Due to the small test amount the significance level was not corrected for multiple testing. All kits and enzymes were used according to the manufacturer’s instructions.

### Purification of XylR

BL21(DE3) cells were transformed with the pET-15b derived constructs for overexpression and purification of His-tagged XylR proteins as previously described with minor modifications [27]. Cells were grown in 50 mL Luria broth at 37 °C and protein expression was induced with 0.5 mM isopropyl-b-D-thiogalactoside when the culture density reached 0.27 g (dcw) L^−1^ at 15 °C. After incubating for 16 h, all cells were harvested (6,750 g for 5 min, 4 °C), washed and resuspended in 1 mL cold lysis buffer (20 mM Tris-HCl pH 8, 300 mM NaCl, 10 mM imidazole). Cells were lysed using a using a FastPrep^®^-24 disrupter. After clarification at 30,000 g (30 min, 4 °C), supernatants were mixed and incubated with 250 μL Ni-NTA agarose (Qiagen) for 1 h at 4. The Ni-NTA agarose containing His-tagged XylR proteins were loaded onto a column, washed with wash buffer (20 mM Tris-HCl pH 8, 300 mM NaCl, 50 mM imidazole) and eluted with elution buffer (20 mM Tris-HCl pH 8, 300 mM NaCl, 300 mM imidazole). PD-10 Desalting Column (GE Health Life Sciences) was used for buffer exchange and imidazole removal. XylR proteins were eluted in 20 mM Tris-HCl pH 8 300 mM NaCl and were quantified using the BCA protein assay kit (Thermo Scientific, Rockford, IL). A single band was observed in a sodium dodecyl sulfate-polyacrylamide gel.

### Electrophoretic Mobility Shift Assay (EMSA)

200 bp DNA fragments containing one XylR binding site (IA and IF) were amplified using *E. coli* XW043 genomic DNA as a template by PCR and purified. EMSA was performed as previously described [42] with minor modifications. A 10 μL reaction mixture (75 mM NaCl, 20 mM Tris-HCl pH 7.5, 1 mM dithiothreitol, 0.1 mg mL^−1^ bovine serum albumin) contained 1.5 nM DNA fragments, 2 mM xylose and varied concentrations of XylR variants ranging from 0 to 400 μM. To test the effect of xylose concentrations on DNA binding (IA site used as a representative), the concentration of XylR and its variants were fixed to 100 nM and xylose concentrations ranging from 0 to 100 μM were employed. After incubation for 15 min at 30 *°C* Orange DNA Loading Dye from Thermo Scientific was added to the reaction mixture to a final 0.5-fold concentration. Bound and unbound DNA fragments were separated by electrophoresis using a native polyacrylamide gel (Mini-PROTEAN^®^TGX^™^ Precast Gel from Bio-Rad) at 5 V cm^−1^ for 3 h at 4 °C. Gels were stained with 40 pg mL^−1^ ethidium bromide and destained with water. Pixel densities were determined with ImageJ and fitted to the non-linear regression model 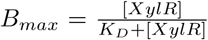 with *B_max_* as the maximum fraction of bound DNA and *K_D_* as the dissociation constant using R.

---

**Figure S1:**
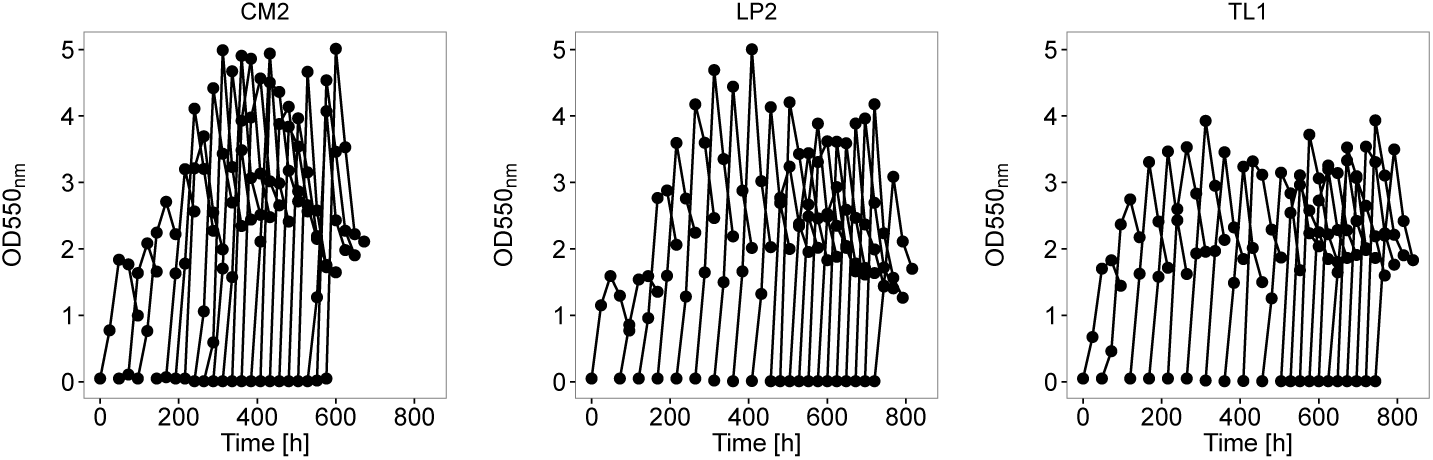
Growth during adaptive laboratory evolution. Optical density at 550 nm was determined every 24 h for the populations that yielded the strains CM2, LP2, and TL1.

**Figure S2:**
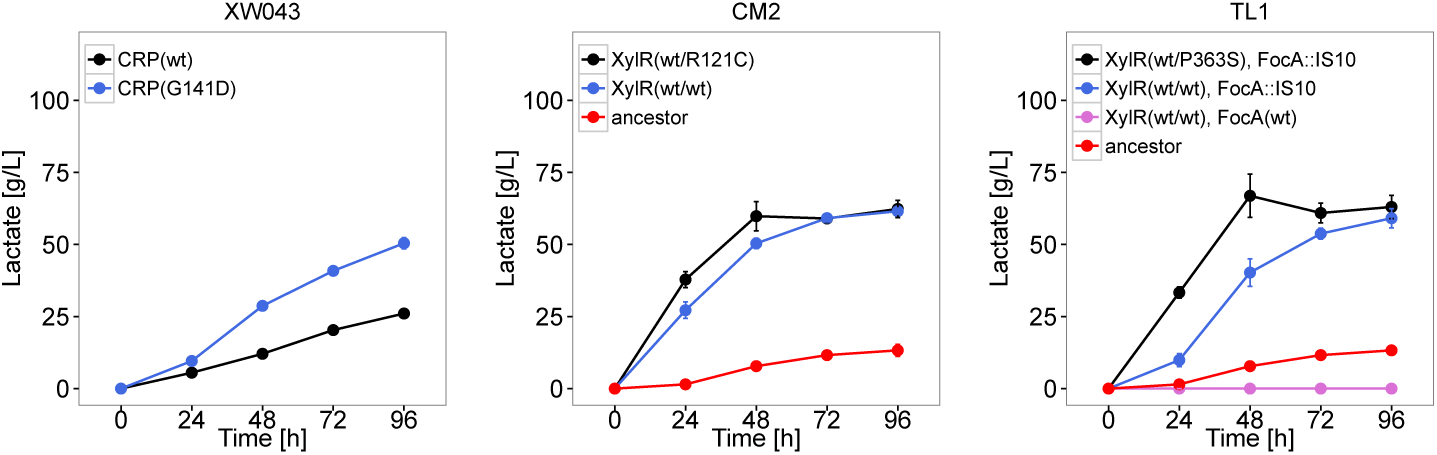
D-lactate production influenced by different genotypes for *crp, focA*, and *xylR*. Through allelic replacements, the genotypes of appropriate genes were modified in different genetic backgrounds: the ancestral XW043, the evolved CM2 and TL1. Lactate concentrations in fermentation broth were measured every 24 h for a 96 h fermentation. Point mutations are indicated by the relevant amino acid substitution. Error bars represent SEM with *n* = 3. wt, wild-type.

**Figure S3:**
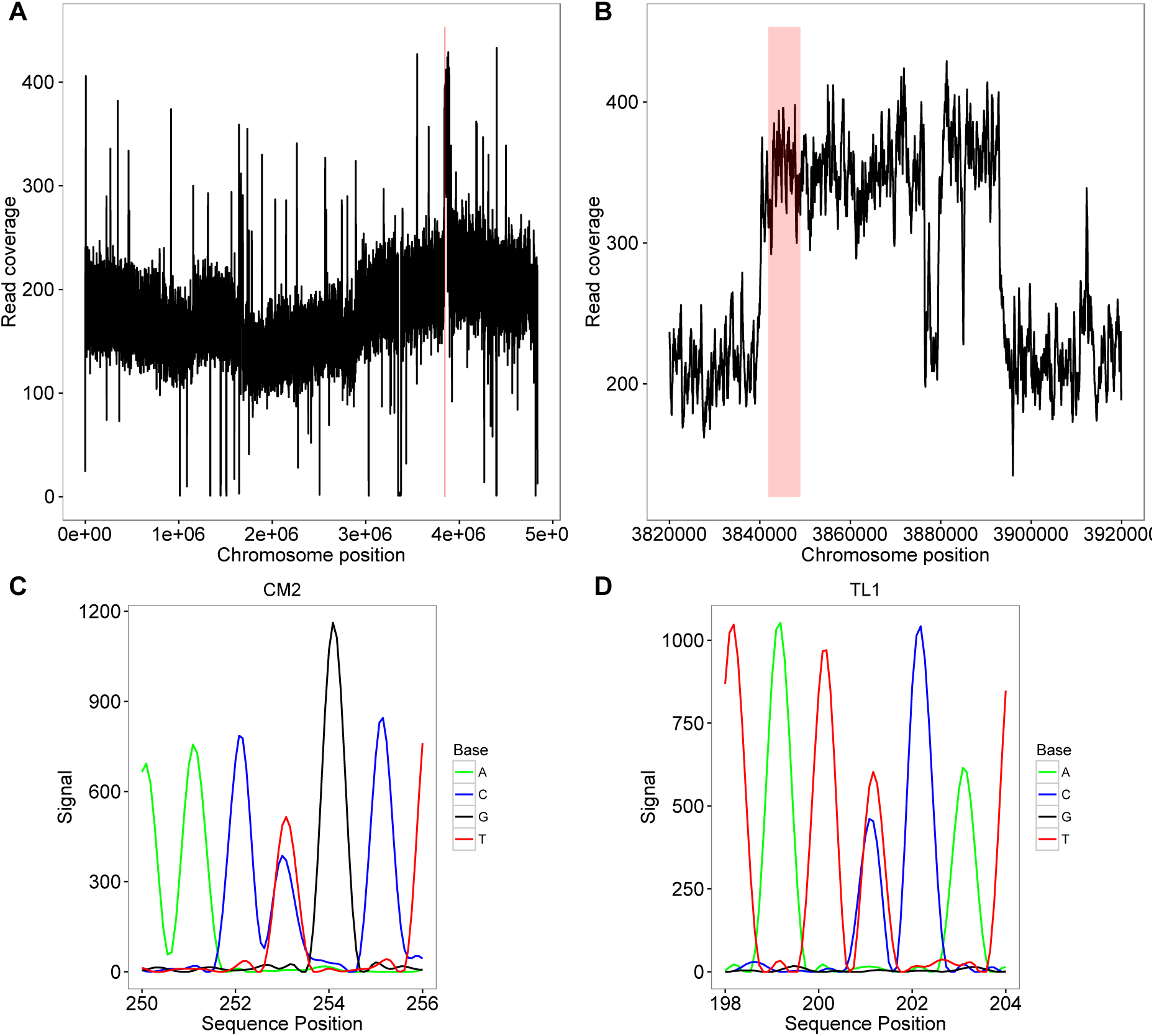
The chromosomal region containing *xylR* is duplicated in XW043. Reads were mapped against the genome of LY180. Read coverage of the whole genome (A) and a close-up (B). The higher coverage compared to the mean indicates duplication. The region encoding *xylAB, xylFGH*, and *xylR* is highlighted in red. Sanger sequencing results for *xylR* in CM2 (C) and TL1 (D) exhibit double peaks at one site and indicate quasi-heterozygous genotypes for *xylR* in both strains.

**Figure S4:**
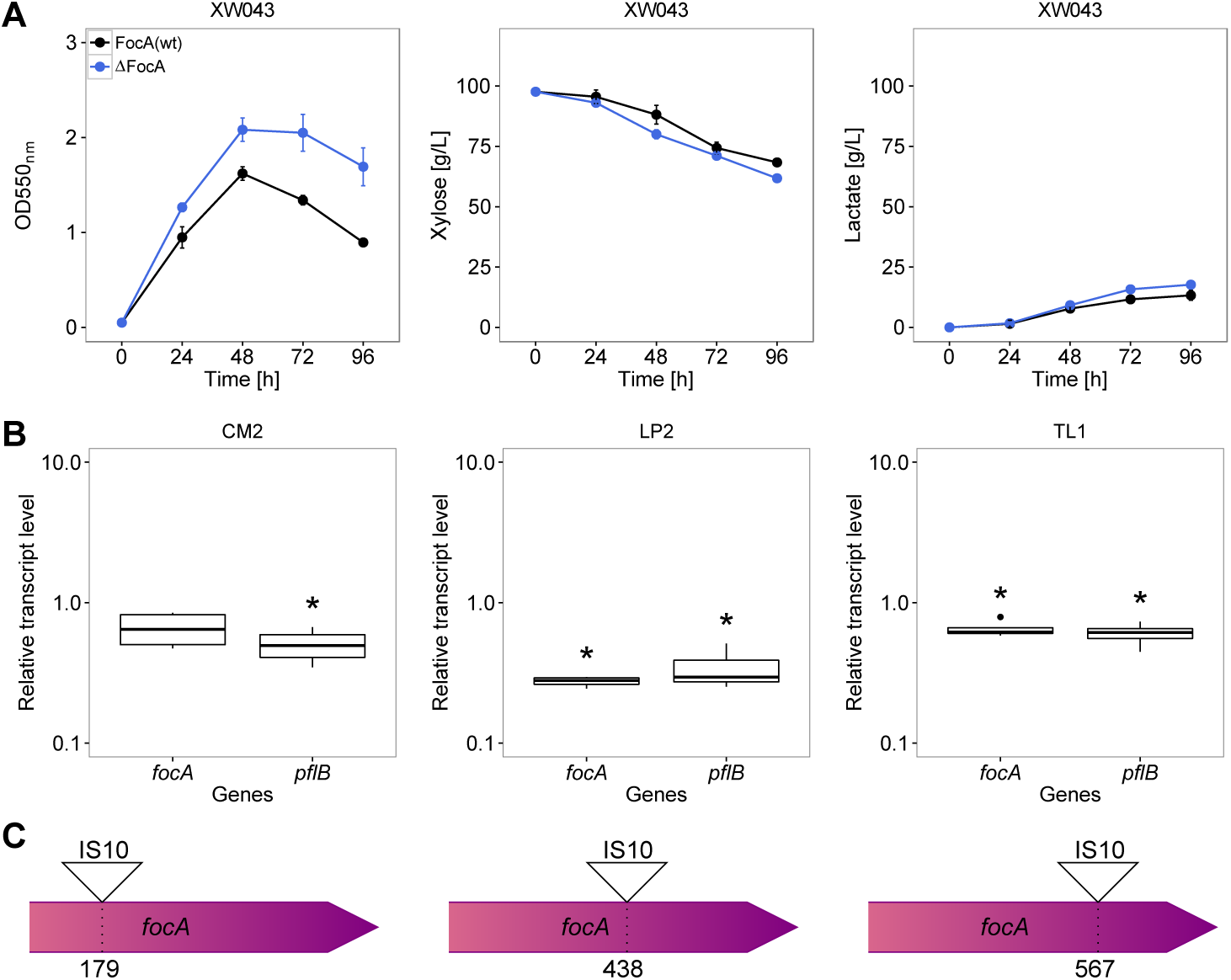
Characterization of the *focA* mutation with respect to lactate fermentation. (A) Fermentation of XW043 and the modified strain with scarless chromosomal *focA* deletion. Extracellular xylose, lactate, and OD550_nm_ were determined. Error bars represent SEM with *n* = 3. wt, wild-type. (B) Relative transcript levels for *focA* and *pflB* of CM2, LP2, and TL1 clones, respectively. The data indicate the fold change of transcript level in relation to the ancestral XW043 background and is normalized by the transcript level of 16S ribosomal RNA (*rrsA*), *n* = 4, unpaired Student’s t-test indicates significance at *p* < 0.05 (*). (C) Schematic drawing of the IS10 insertion in the *focA* reading frame of CM2, LP2, and TL1. The numbers indicate the determined insertion position relative to the translation start point.

**Figure S5:**
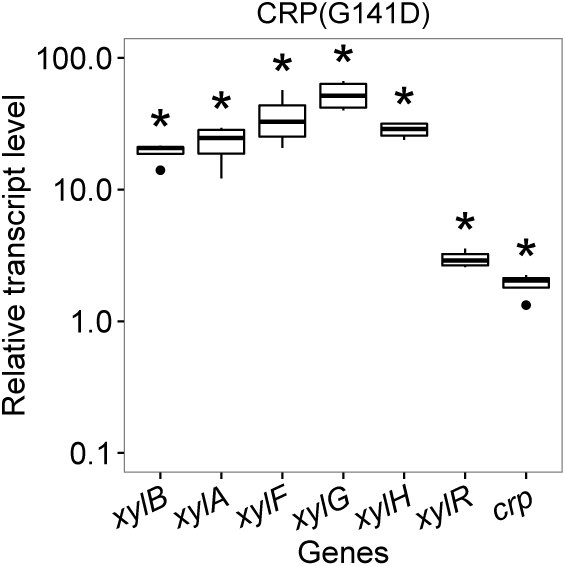
Relative transcript level of selected genes of LP2 with the CRP variant G141D. The data indicates the fold change of transcript level in relation to the ancestral XW043 background and is normalized by the transcript level of 16S ribosomal RNA (*rrsA*), *n* = 4, unpaired Student’s t-test indicates significance at *p* < 0.05 (*).

**Figure S6:**
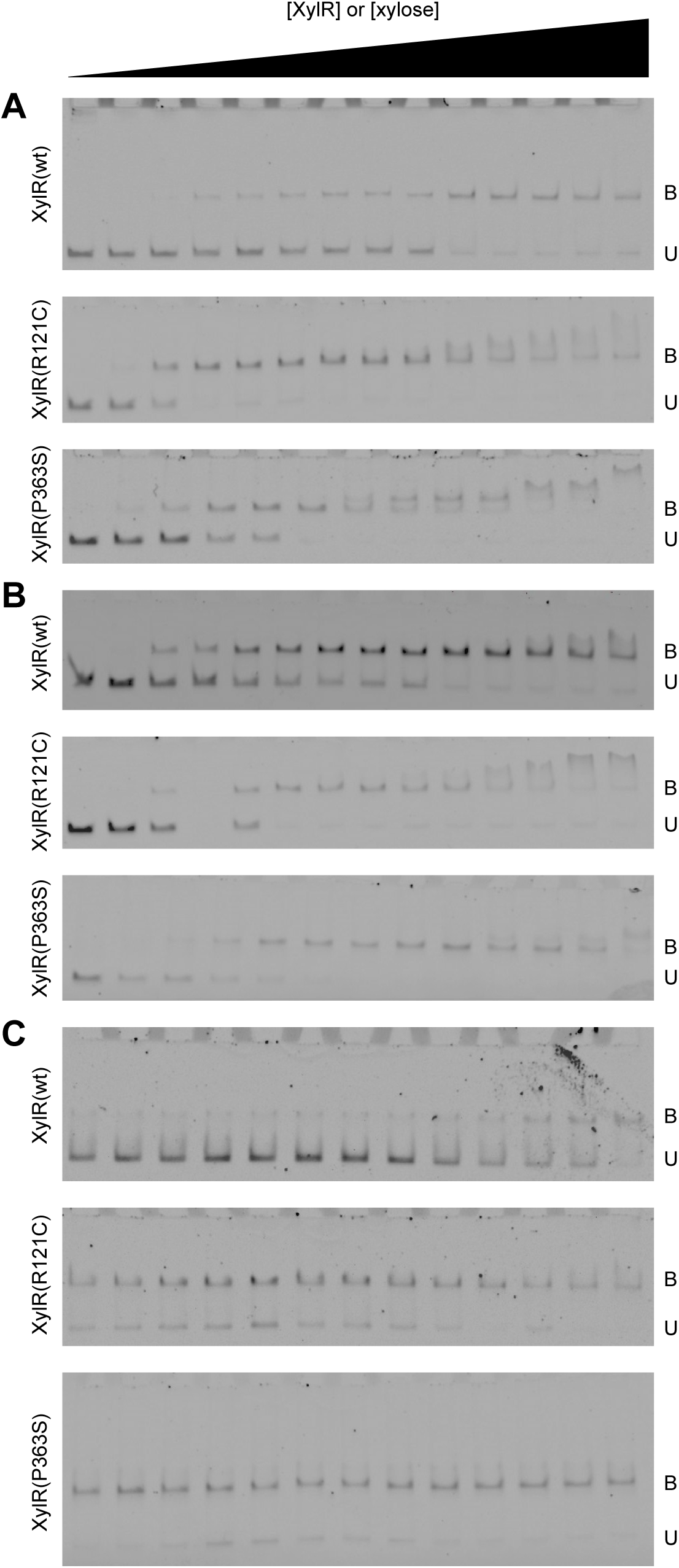
Electrophoretic mobility shift assays testing the binding affinity of XylR variants. XylR wild-type (wt) and the variants R121C and P363S were incubated with DNA containing the known binding sites IA (A) and IF (B) with increased protein concentrations from 0 to 400 nM in the presence of 2 mM xylose. To test the effect of xylose concentrations (C), xylose concentrations ranging from 0 to 100 pM were employed in the presence of 100 μM different XylR variants. Bound (B) and unbound (U) fraction of DNA fragments are indicated.

**Figure S7:**
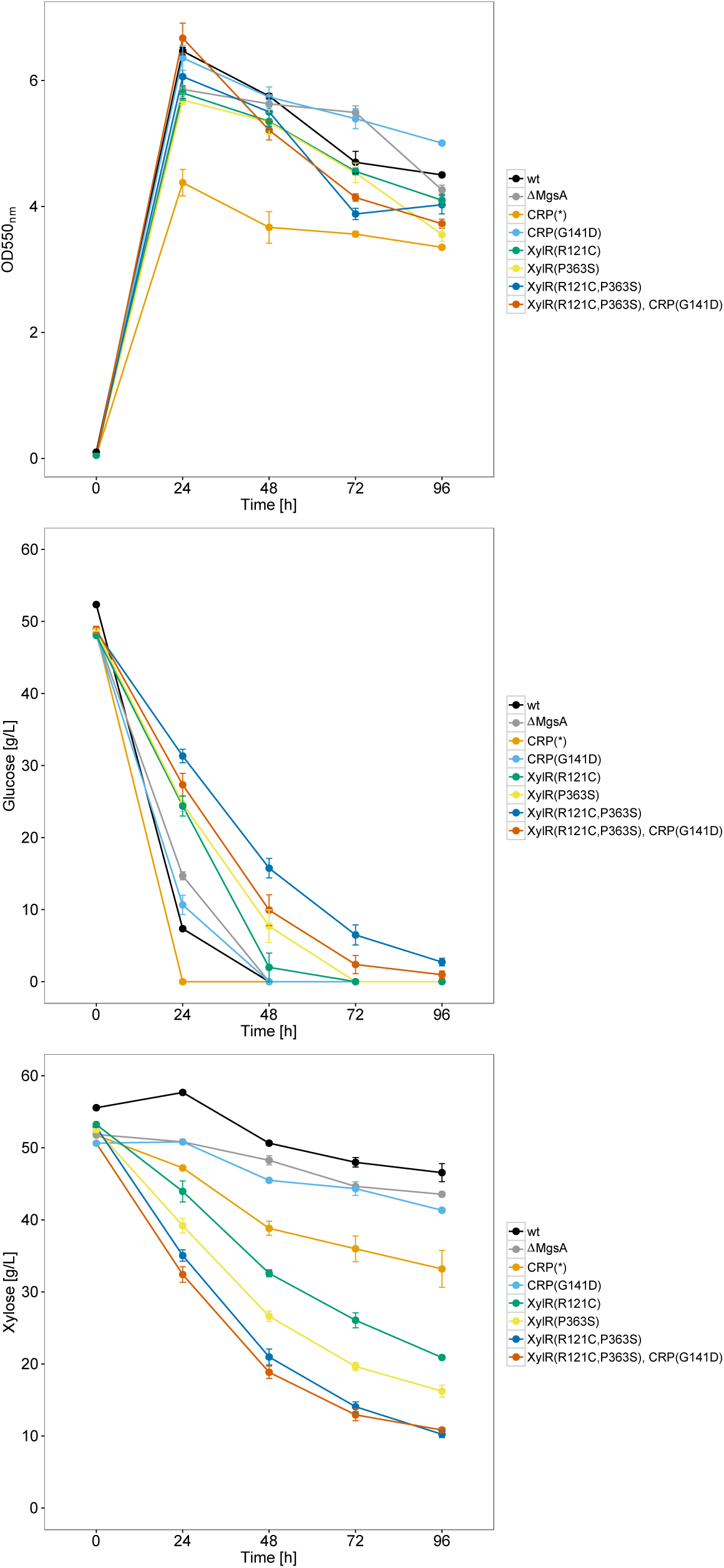
Co-utilization of glucose-xylose mixtures by batch fermentations. *E. coli* W was engineered with different genotypes including previously reported genetic traits such as *mgsA* deletion (ΔMgsA), CRP(*) and identified CRP and XylR variants to compare their effect of releasing CCR. Batch fermentations were performed using glucose-xylose mixtures (50 g L^−1^ of each sugar) for 96 h of fermentation. Cell optical density (0D550_nm_), extracellular glucose and xylose concentrations were determined. Error bars represent SEM with *n* = 3. wt, wild-type.

**Table S1:**
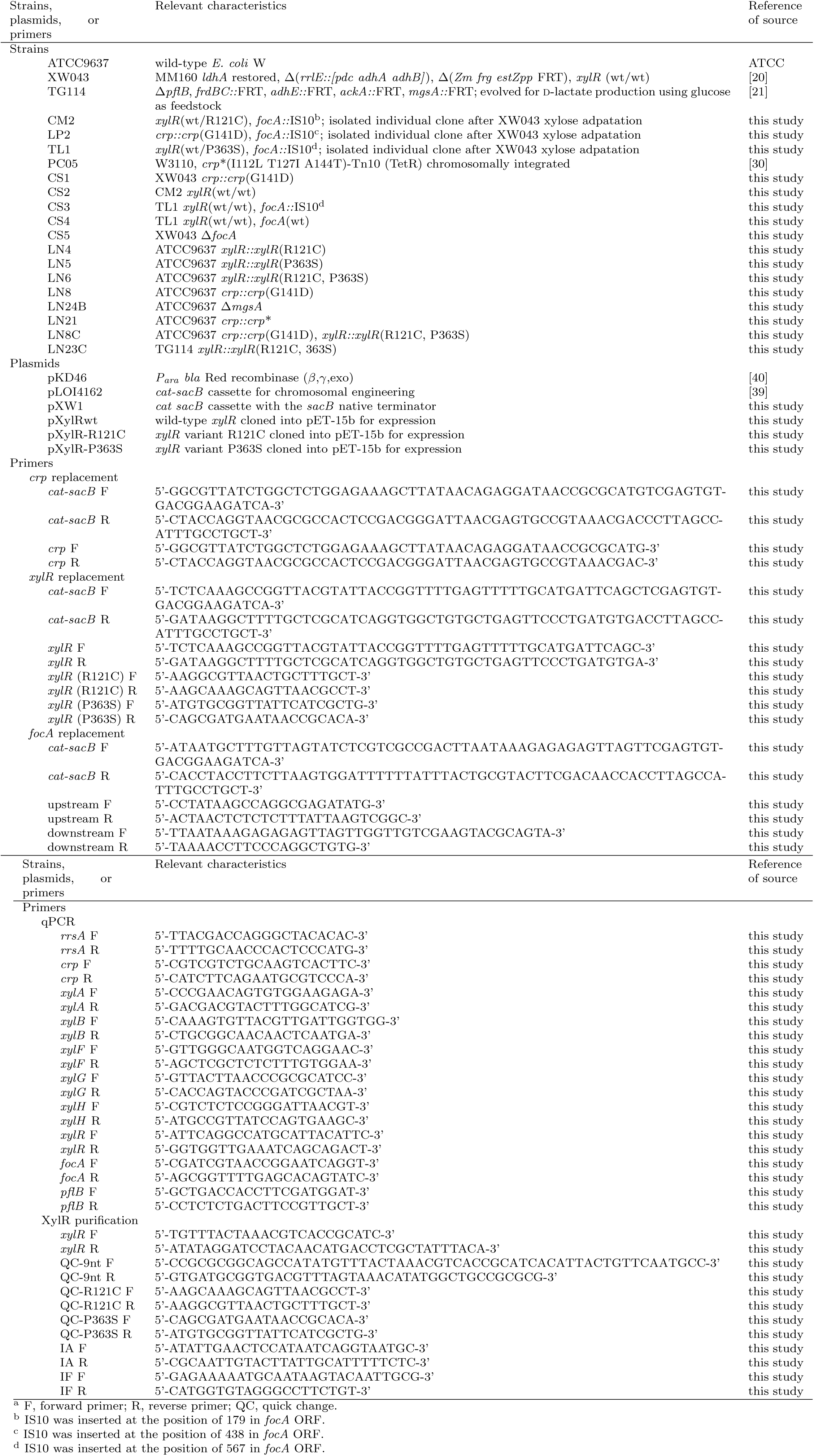
Strains, plasmids, and primers used in this study^a^.

**Table S2:**
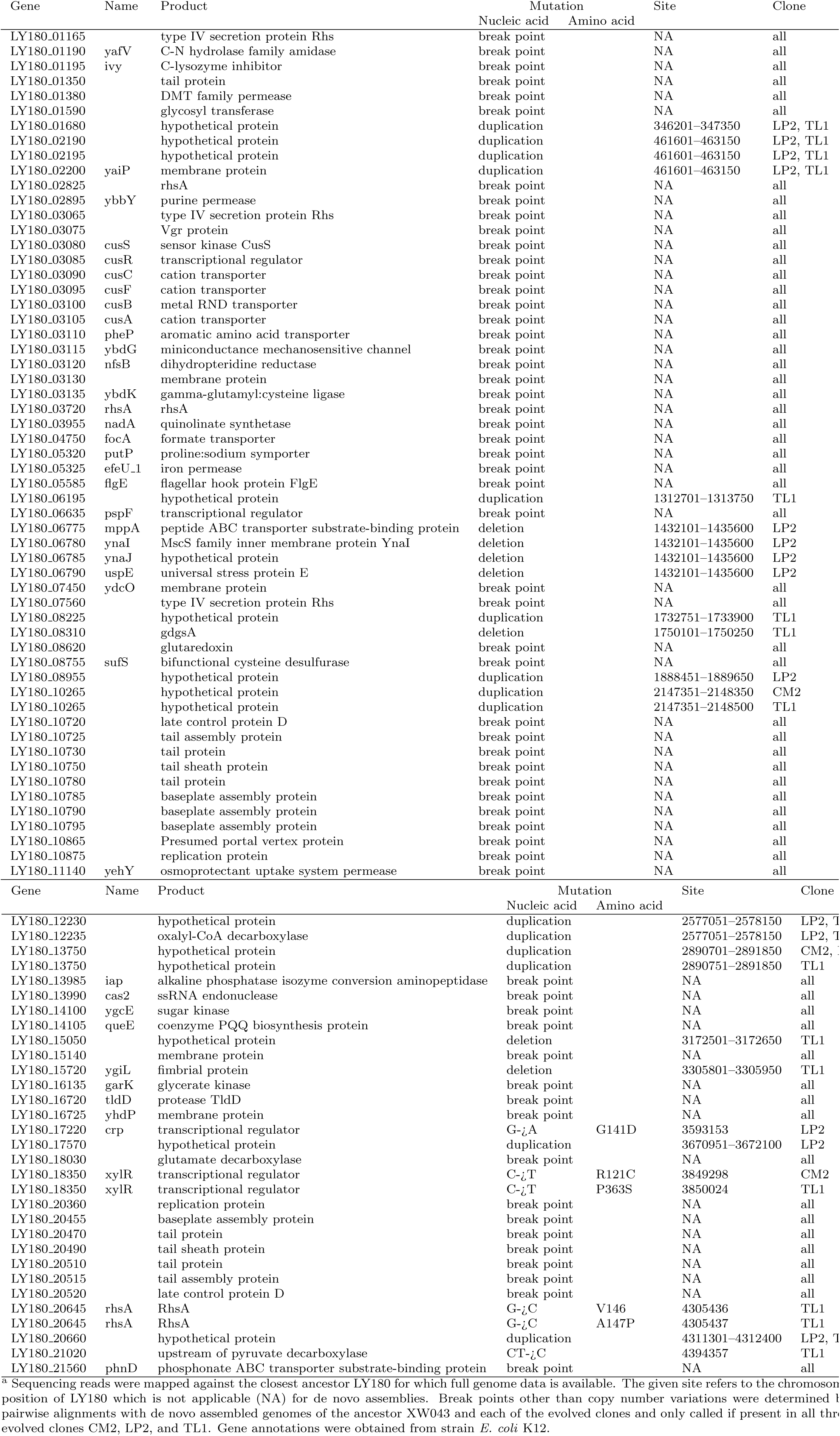
Identified SNPs, copy number variations, and miscellaneous break points unique to LP2, CM2, and TL1 compared to ancestor strain XW043^a^.

